# AnkG-Neurofascin complex structure reveals binding mechanisms required for integrity of the AIS

**DOI:** 10.1101/2022.04.27.489743

**Authors:** Liping He, Wenli Jiang, Jianchao Li, Chao Wang

## Abstract

The axon initial segment (AIS) has characteristically clustering of voltage-gated sodium channels (Nav), cell adhesion molecule Neurofascin (Nfasc), and neuronal scaffold protein Ankyrin-G (AnkG) in neurons, which facilitate generation of action potential and maintenance of axonal polarity. However, the mechanisms underlying AIS assembly and maintenance remain poorly understood. Here we report the high-resolution crystal structure of the AnkG in complex with a fragment from Nfasc cytoplasmic tail that shows, in conjunction with binding affinity assays, the molecular basis of AnkG-Nfasc binding. We confirm AnkG interacts with the FIGQY motif in Nfasc, and identify another region required for their high affinity binding. Structural analysis revealed that ANK repeats form four hydrophobic or hydrophilic layers in the AnkG inner groove that coordinate interactions with Nfasc. Moreover, disruption of the AnkG-Nfasc complex abolishes Nfasc enrichment at the AIS in hippocampal neurons. Finally, structural and biochemical analysis indicated that L1 syndrome-associated mutations in L1CAM compromise binding with ankyrins. These results define the mechanisms underlying AnkG-Nfasc complex formation and show that AnkG-dependent clustering of Nfasc is required for AIS integrity.

## Introduction

Neurons are highly polarized cells typically composed of a soma, multiple dendrites, and a single axon. The specialized compartmentalization of neurons is critical for their physiological functions. The axon initial segment (AIS) connects the soma and the axon, serving as a vital region responsible for initiating action potential and maintaining neuronal polarity (1-3). Several studies conducted over the past decades have expanded our understanding of the molecular architecture and organization of the AIS (4-7). The AIS is characterized by highly dense enrichment with a variety of proteins that include ion channels, cell adhesion molecules, scaffold proteins, regulatory proteins, and cytoskeletal proteins (8-10). Among these proteins, Ankyrin-G (AnkG) is considered as the master organizer that directs the recruitment of diverse components to the AIS (4, 5). Previous studies have demonstrated that AnkG interacts with Nav1.2, a primary ion channel involved in the initiation of action potential (11, 12), as well as with Ndel1, a dynein regulator that functions in selective sorting and polarity maintenance at the AIS (13, 14). In addition, AnkG is also well-known to interact with Neurofascin 186 (Nfasc), a cell adhesion molecule that is essential for AIS integrity (15, 16). Loss of AnkG leads to disruption of axonal polarity and disassembly of the AIS, either *in vivo* or *in vitro* (3, 17).

In humans, the ankyrin family contains three known members: AnkR, AnkB and AnkG, which are all ubiquitously expressed and perform non-redundant functions in most tissues (18, 19). In neurons, the AIS and nodes of Ranvier are specifically enriched with 480/270 kDa AnkG alternative splice variants (20, 21). Clustering of AIS membrane proteins, including sodium channels, potassium channels, Neurofascin 186, and NrCAM has been proposed to depend on their specific interactions with the ankyrin repeats (ANK repeats) of AnkG (15, 22, 23). In earlier studies, by solving the crystal structures of ANK repeats in complex with its own autoinhibitory segments or Nav1.2, we proposed that ankyrins can utilize a combination of multiple binding sites in the inner groove of its ANK repeats to interact with various membrane protein targets (*e*.*g*., Nav1.2, Nfasc) (24, 25). However, more high-resolution structures of ankyrins in complex with their targets are needed to fully disclosure the diverse target recognition mechanisms of ANK repeats.

Neurofascin 186 is a membrane-spanning cell adhesion molecule belonging to the L1 group of the immunoglobulin superfamily (L1 family) proteins, that include L1CAM, Neurofascin, NrCAM, and CHL1 (26, 27). Neurofascin 186 is a major splicing variant that is predominantly expressed in neurons, and is restricted to the AIS and nodes of Ranvier (28). Loss of Nfasc leads to dissociation of the AIS in Nfasc-null mice (29). A recent study showed that Nfasc is highly mobile after its recruitment to the axonal membrane and diffuses bidirectionally until anchored at the AIS through its interaction with AnkG (30). These findings thus highlight the importance of the Nfasc-AnkG complex in forming the AIS. Previous studies have suggested that Nfasc targeting to the AIS is dependent upon AnkG binding through a FIGQY motif in its cytoplasmic domain (31, 32). However, despite a wealth of available functional data, the mechanistic details and structural basis of their interaction remain poorly understood.

In the present study, we characterized the binding and structural interactions between AnkG and Nfasc in detail. Analysis of truncation variants showed that a segment (1187-1214) in the cytoplasmic tail of Nfasc mediates its strong interaction with AnkG. We then elucidated the precise molecular mechanisms governing this interaction by solving the crystal structure of the ankyrin-binding region of Nfasc in complex with the ANK repeats of AnkG. In addition to the recognized FIGQY motif, our structural data identified an N-terminal region that is also essential for binding. Further structural analysis highlighted a pattern of preferential distribution of residues in four hydrophobic or hydrophilic layers formed by ANK repeats in the inner groove responsible for targets binding. Moreover, we showed that disruption of AnkG binding abolishes Nfasc localization to the AIS in primary cultured hippocampal neurons. We also confirmed that AnkG can interact with all four members of the L1 family and found that, two L1 syndrome-associated mutations in L1CAM interfere with its binding to ankyrins. These results provide inroads to understanding the possible pathological mechanisms underlying ankyrin-L1 family complex-related neuronal diseases.

## Results

### Nfasc ABD in the cytoplasmic tail mediates binding with AnkG

Previous studies have established that Nfasc binds to the AnkG ankyrin repeats (ANK repeats) region through interaction with its C-terminal cytoplasmic tail (15, 31). To investigate the biochemical details of this interaction, we used purified AnkG ANK repeats (residues 38-855, hereafter R1-24) and two fragments of Nfasc (including the full-length Nfasc cytoplasmic tail, residues 1132-1240, and the ankyrin binding domain, ABD, residues 1187-1214) for fast protein liquid chromatography (FPLC) and isothermal titration calorimetry (ITC) assays to evaluate the binding affinity and map the specific sites that participate in their interaction (Fig. 1A, B). Both FPLC and ITC results showed that the Nfasc ABD interacts strongly with AnkG R1-24 (Fig. 1C, D). Furthermore, Nfasc ABD is both necessary and sufficient for binding to AnkG with a dissociation constant (K_d_) of ∼ 0.22 μM (Fig. 1G, the first two rows and the last row, & Fig. S1).

**Fig. 1.**
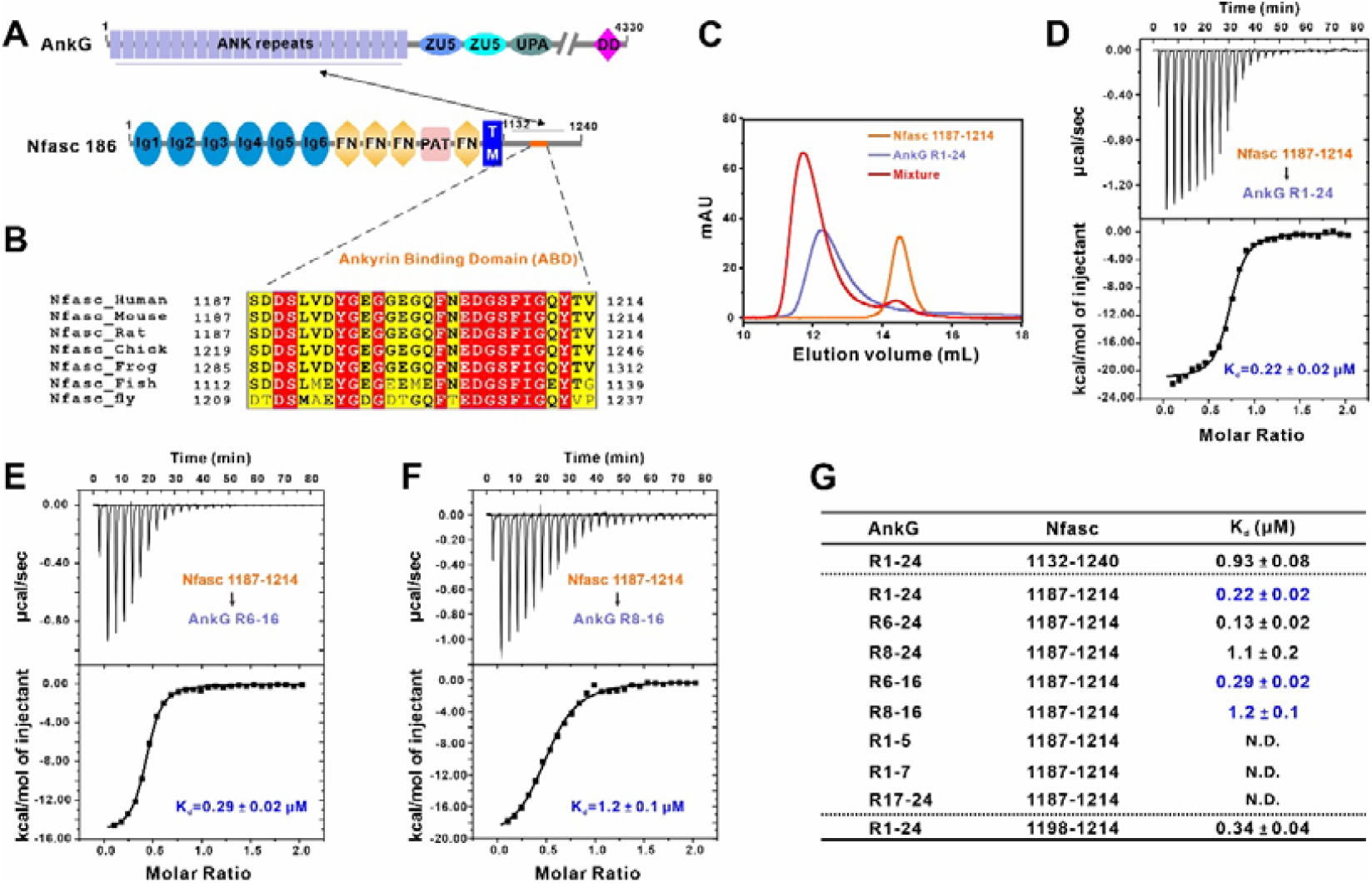
Nfasc ABD in the cytoplasmic tail mediates binding with AnkG. (A) Schematic diagram showing the domain organizations of AnkG and Nfasc. In this drawing, the interaction between AnkG and Nfasc is indicated by a 2-way arrow. The ankyrin binding domain (ABD) of Nfasc are highlighted in orange in the cytoplasmic region. DD, death domain. (B) Sequence alignment of Nfasc ABD in different species showing a high conservation through evolution. Residues that are identical and highly similar are indicated in read and yeallw boxes, respectively. (C) Analytical gel filtration analysis showing that Nfasc ABD (residues 1187-1214) and AnkG ANK repeats (R1-24, residues 38-855) interacted with each other. (D-F) ITC-based measurements of the binding affinity of Nfasc 1187-1214 with AnkG R1-24 (D), AnkG R6-16 (E), or AnkG R8-16 (F). The K_d_ error is the fitting error obtained using one site binding kinetics model in Origin 7.0 to fit the ITC data. (G) The measured binding affinities between different Nfasc fragments and various truncations of AnkG ANK repeats from ITC-based binding assays. N.D. indicates that no binding was detected.

Based on findings established in our previous studies of ANK repeats (24), we designed several AnkG truncation variants to identify the precise Nfasc binding site. Through comprehensive analyses by ITC and FPLC, we found that the major binding region was located between the eighth and sixteenth ANK repeats (AnkG R8-16), while the first seven repeats and last eight repeats exhibited no binding activity with Nfasc (Fig. 1E-G & Fig. S1 & Fig. S2). The binding affinity between the AnkG R6-16 and Nfasc showed a stronger trend than that of AnkG R8-16 (K_d_ ∼0.29 μM vs K_d_ ∼1.2 μM), which suggested that the two N-terminal repeats could possibly involve in binding (Fig. 1E-G). Taken together, these results showed that Nfasc interacts with AnkG strongly and the ABD of the Nfasc cytoplasmic tail mediated its interactions with AnkG.

### Overall crystal structure of AnkG in complex with Nfasc ABD

In order to better understand the molecular mechanisms governing AnkG-Nfasc complex formation, we sought to solve the crystal structure of AnkG-Nfasc in complex. To this end, we tested various preparations of the protein complex or combinations of fusion proteins guided by the results of our above binding assays. After extensive efforts, we successfully obtained crystals of Nfasc ABD fused with the AnkG R8-16 region that diffracted to 2.5 Å resolution and we subsequently solved the complex structure using molecular replacement (Fig. 2 & Table S1).

**Fig. 2.**
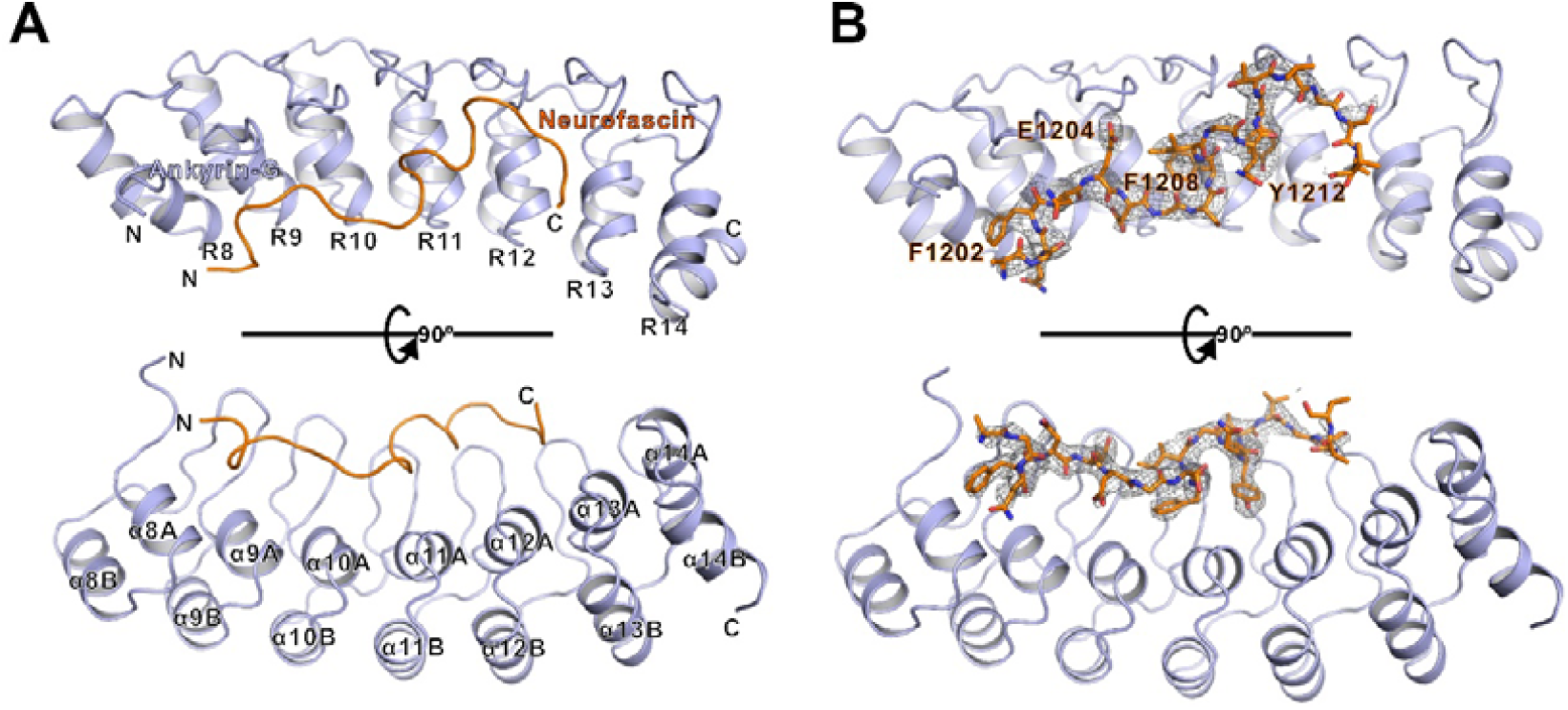
Overall crystal structure of AnkG in complex with Nfasc ABD. (A) Ribbon representation model showing the overall structure of the AnkG-Nfasc complex. In this drawing, AnkG is shown in light blue, and Nfasc is shown in orange. Nfasc peptide extends in the inner groove of ANK repeats with parallel orientation. The coordinates and structure factors of the AnkG and Nfasc complex have been deposited to the Protein Data Bank under the accession code number 7XCE. (B) Electron density map of Nfasc in the ribbon diagram of AnkG-Nfasc complex with critical residues including F1202, E1204, F1208, and Y1212 highlighted.

In this complex, the AnkG R8-14 adopted a canonical ANK repeat architecture (Fig. 2A, B), although it should be noted that the last two ANK repeats (R15 and R16) were missing in the final structure. In agreement with our understanding of ankyrin structure, the Nfasc ABD peptide extended into the inner groove formed by the ANK repeats. However, in contrast with the antiparallel orientation (N-terminal to C-terminal binding) observed in other ANK repeats-target complex structures reported for ankyrin family proteins (Nav1.2-AnkB, AnkR-AnkB, etc) (24), the AnkG-Nfasc complex adopted an unexpected parallel binding orientation (*i*.*e*., N-terminal to N-terminal binding between Nfasc and ANK repeats). Although we used the Nfasc 1187-1214 segment for crystallization, only the Nfasc 1198-1214 region could be clearly traced in the electron density map (Fig. 2B). Furthermore, this 1198-1214 region retained the majority of its binding ability with AnkG, which was in line with our findings in truncation variants (Fig. 1G). Given the parallel orientation combined with the above-mentioned biochemical data, it was reasonable to speculate that the Nfasc 1187-1197 residues might facilitate binding through interaction with the N-terminal ANK repeats adjacent to R8-16 in AnkG.

### Hydrophobic and hydrogen bonding interactions at the AnkG-Nfasc interface

We next examined the specific interactions between residues responsible for their binding and found that the AnkG-Nfasc interface is mainly mediated by hydrophobic and hydrogen bonding interactions. In particular, F1202 from Nfasc inserts into a hydrophobic pocket formed by I277, V282, and C315 from AnkG (Fig 3A & Fig. 4A). Similarly, F1208, another aromatic residue from Nfasc, occupies the hydrophobic groove formed by AnkG residues L343, L376, and V381 (Fig 3A & Fig. 4A). In addition to these central hydrophobic interactions, several hydrogen bonds also contribute to the high affinity and specificity of the interaction. Among these, we found that the E1204 sidechain from Nfasc forms hydrogen bonds with T339 and N341 in AnkG, while the main chain of E1204 forms another hydrogen bond with the sidechain of D308 from AnkG (Fig. 3A). In addition, D1205 from Nfasc forms hydrogen bonds with R318 and Q351 from AnkG (Fig. 3A). It should be noted that several other hydrogen bonding pairs, including T306^AnkG^-N1203^Nfasc^, H384^AnkG^-S1207/Y1212^Nfasc^, and D374^AnkG^-G1210/T1213^Nfasc^, also contribute to stabilizing the complex assembly (Fig. 3A). Collectively, ∼989 Å^2^ total surface area (calculated by PISA server, https://www.ebi.ac.uk/pdbe/pisa/) participated in the interaction between the AnkG ANK repeats and the Nfasc ABD peptide.

**Fig. 3.**
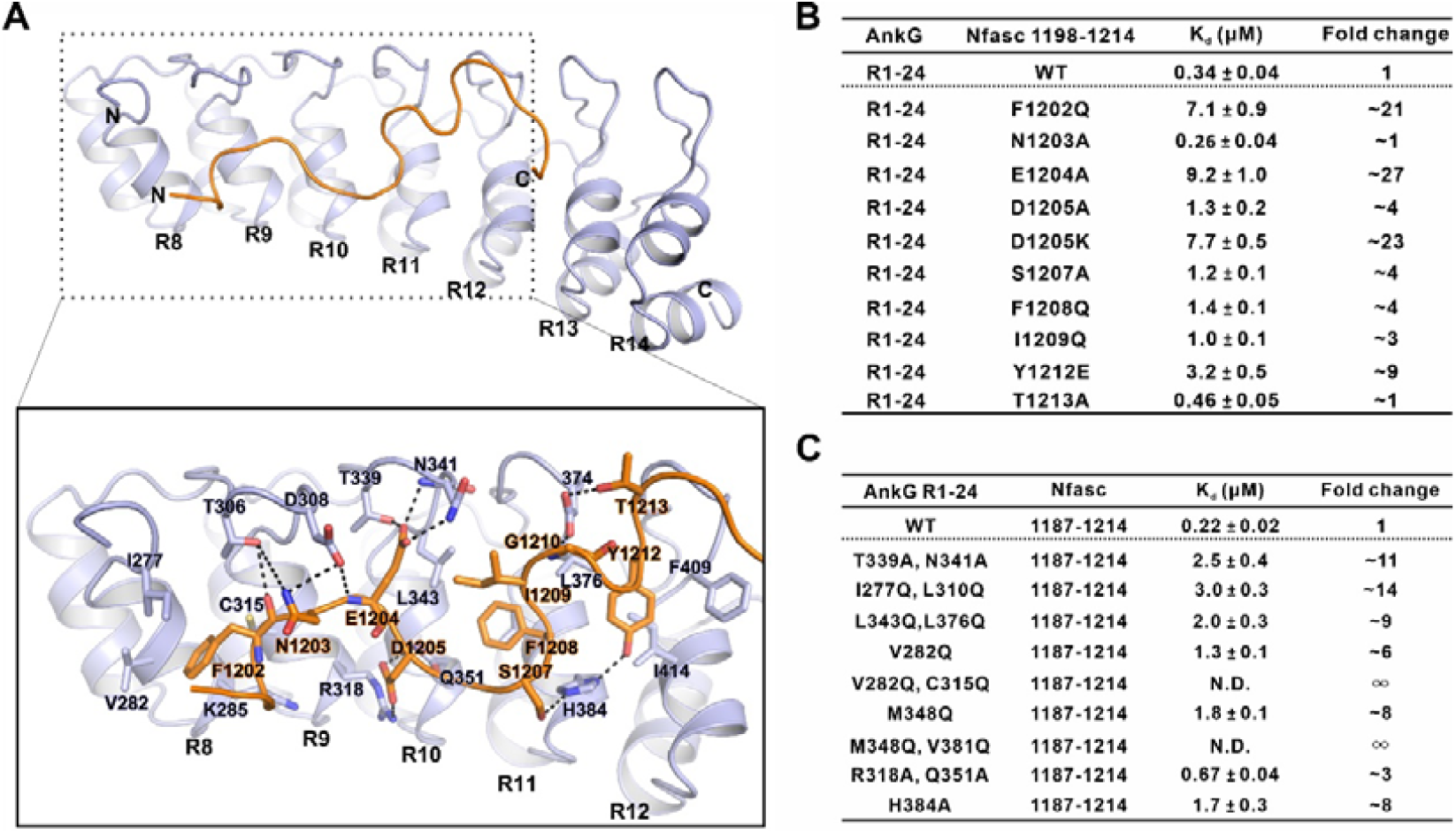
The interface of AnkG-Nfasc complex. (A) Ribbon diagram showing the detailed interactions between AnkG and Nfasc. Residues involved in the binding are shown in stick model. The hydrogen bonds are shown as dotted lines. (B) The measured binding affinities between various mutations of Nfasc 1198-1214 and AnkG R1-24 based on ITC assays and comparison against Nfasc 1198-1214 WT (wild-type) in binding with AnkG R1-24. (C) The measured binding affinities between various mutations in each layer of AnkG R1-24 and the Nfasc 1187-1214 based on ITC assays and comparison against AnkG R1-24 WT in binding with Nfasc 1187-1214.

**Fig. 4.**
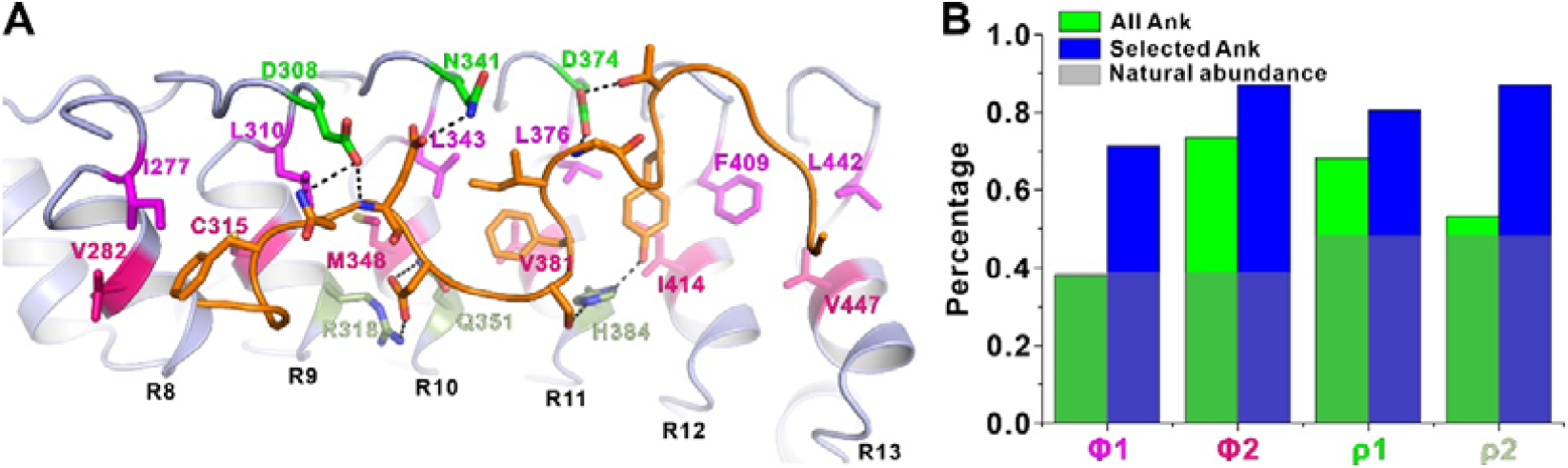
Amino acids in ANK repeats show distribution preference in inner groove-dependent binding. (A) The amino acids layers of ANK repeats in the interface of AnkG-Nfasc complex. Amino acids of four layers in ANK repeats displayed patterned distributions are highlighted in stick model. Layer 1 (green) and layer 4 (pale green) are polar residues while layer 2 (magenta) and layer 3 (hot pink) are hydrophobic residues. The hydrogen bonds are shown as dotted lines. (B) Histogram of residues distribution preference analysis of the ANK repeats in inner groove-dependent binding. Green boxes represent all ANK repeats. Blue boxes represent the ANK repeats that adopt the inner groove-dependent binding pattern. Gray boxes represent the natural distribution of the residues in all human proteins. Φ1, Φ2, ρ1, and ρ2 are corresponding to layer 2, layer 3, layer 1, and layer 4 positions, respectively. The vertical axis represents the percentage of hydrophobic residues (Φ1, Φ2) or polar residues (ρ1, ρ2) in ANK repeats.

To further confirm the interaction mode between AnkG and Nfasc, we evaluated the impact of specific mutations in either AnkG or Nfasc at the binding interface. In line with our structural analysis, ITC-based assays showed that mutations in the Nfasc ABD which disrupted hydrophobic interactions generally weakened AnkG-Nfasc interaction (Fig. 3B). For example, an F1202Q conversion mutation in the Nfasc ABD variant partially disrupted hydrophobic interactions and led to a greater than 20-fold decrease in binding (Fig. 3B & Fig. S3A). Single substitutions of Gln at the F1208 or I1209 residues in Nfasc led to a slight decrease of their interaction (Fig. 3B & Fig. S3G, H), whereas an Ala substitution at E1204 in Nfasc resulted in a significant, ∼27-fold decrease in binding affinity (Fig. 3B & Fig. S3C), and D1205A/K substitutions in Nfasc variants also weakened its binding with AnkG (Fig. 3B & Fig. S3D, E). These findings thus illustrated the pivotal role of these hydrogen bonds at the ABD-ANK repeat interface in the crystal structure. Notably, Nfasc residue Y1212, which is a likely phosphorylation site, was reported to act as a molecular switch for regulating the interaction between Nfasc and AnkG (32). However, we found that a Y1212E phosphorylation mimic variant exhibited only an approximately 9-fold decrease in binding, potentially due to incomplete imitation of the *in vivo* effects of phosphorylation (Fig. 3B & Fig. S3I). Overall, the observed hydrophobic and hydrogen bonding/electrostatic interactions between AnkG and Nfasc strongly resembled those interactions between the inner groove of ankyrins and short peptides of other targets described in previous studies (24, 25).

### Amino acids in ANK repeats show distribution preference in inner groove-dependent binding

Next, we evaluated the specific roles of AnkG residues required for binding with Nfasc in the inner groove. With great interest, we found that residues in the ANK repeats located in inner groove exhibited clear patterns of distribution to form distinct hydrophilic layers and hydrophobic layers (Fig. 4A). More specifically, polar residues from the upper loop connecting two adjacent ANK repeats (D308, N341 and D374) formed the first, hydrophilic, layer (green). The second, hydrophobic, layer (magenta) consisted of residues from the hairpin region, including I277, L310, L343 L376, F409 and L442. Residues from the belly of the αA (V282, C315, M348, V381, I414 and V447) formed the third, hydrophobic, layer (hot pink), while residues from the bottom of αA (R318, Q351, and H384) formed the fourth, hydrophilic, layer (pale green) (Fig. 4A). Obviously, residues in the hydrophilic layers participated in the electrostatic or hydrogen bond interactions with N1203, E1204, D1205, S1207, Y1212, and T1213 from Nfasc, and residues in the hydrophobic layers were responsible for coordination of hydrophobic residues of Nfasc, including F1202, F1208 and I1209 (Fig. 3A & Fig. 4A).

We then introduced corresponding mutations into each hydrophilic or hydrophobic layer of AnkG and, consistent with these predicted functions, ITC assays showed that all of these mutations could decrease the binding affinity for the Nfasc ABD (Fig. 3C & Fig. S4). Among them, double substitutions for Gln in the hydrophobic residues of the hydrophobic third layer completely abolished binding with the Nfasc ABD, indicating that these residues in the third layer were indispensable for AnkG-Nfasc interactions (Fig. 3C & Fig. S4E, G).

We found that in several ANK repeats containing proteins (AnkR/B/G, KANK1/2, Espin1/Espin like) that bind their respective target peptides in an inner groove-dependent manner, they all utilize these four layers of residues for binding. We then wondered whether there were amino acid preferences in these four layers of ANK repeats for targets binding. In Φ1 and Φ2 (corresponding to layer 2 and 3) positions, we observed an obvious preference for hydrophobic amino acids (A/C/F/I/L/M/V/W/Y) (Fig. 4B & Fig. S5). Particularly, ∼70% of the Φ1 position of these target binding ANK repeats contains a hydrophobic residue (blue bars), whereas the percentages for this position in all ANK repeats in human proteome (green bars) or the natural abundancy of hydrophobic residues (grey bars) are both ∼40%. Similarly, in the ρ1 and ρ2 (corresponding to layers 1 and 4) positions, polar amino acids (D/E/H/K/N/Q/R/S/T) occurred at higher frequencies than they did in all ANK repeats or throughout the whole proteins (Fig. 4B & Fig. S5). Collectively, these patterns of preferential residue distribution within the four layers indicated that ANK repeats may share similar mechanisms for binding to diverse peptides mediated by the inner groove. Thus, we propose that by analyzing the amino acid properties for these four positions of ANK repeats from a protein, we might be able to determine whether this protein can use the inner groove for target recognition.

### Disruption of AnkG binding abolishes Nfasc enrichment at the AIS

Both AnkG and Nfasc are known to specifically localize at the AIS region in neurons in mice and humans (29, 33, 34). Consistent with these previously reported data, we first confirmed that AnkG and Nfasc both showed clear enrichment and co-localization at the AIS in primary cultured hippocampal neurons obtained from C57BL/6 mice (Fig. 5A, D). To investigate the function of AnkG-Nfasc complex in neurons, we suppressed endogenous Nfasc expression using Nfasc-shRNA at day four of *in vitro* in hippocampal neuron culture (DIV 4). At DIV 7, immunofluorescent staining showed that Nfasc signal was substantially reduced in neurons transfected with BFP-Nfasc-shRNA compared to those transfected with the BFP vector (Fig. 5B, C & E). Interestingly, the intensity of AnkG signal did not change between DIV4 and DIV7, which indicated that enrichment for AnkG at the AIS did not depend on Nfasc at this stage. We then overexpressed an HA-Nfasc (*i*.*e*., wild-type, WT) complementation plasmid or an HA-Nfasc variant plasmid harboring either F1202Q or E1204A conversions in the Nfasc-depleted hippocampal neurons. We found that the WT Nfasc expression restored AIS enrichment while the F1210Q and E1204A Nfasc variants exhibited total loss of enrichment at the AIS (Fig. 5F-I), which was in agreement with our above AnkG binding assays. Taken together, these results indicated that the Nfasc localization to the AIS depends on its binding with AnkG, and that disruption of this interaction results in Nfasc failure to localize at the AIS in neurons.

**Fig. 5.**
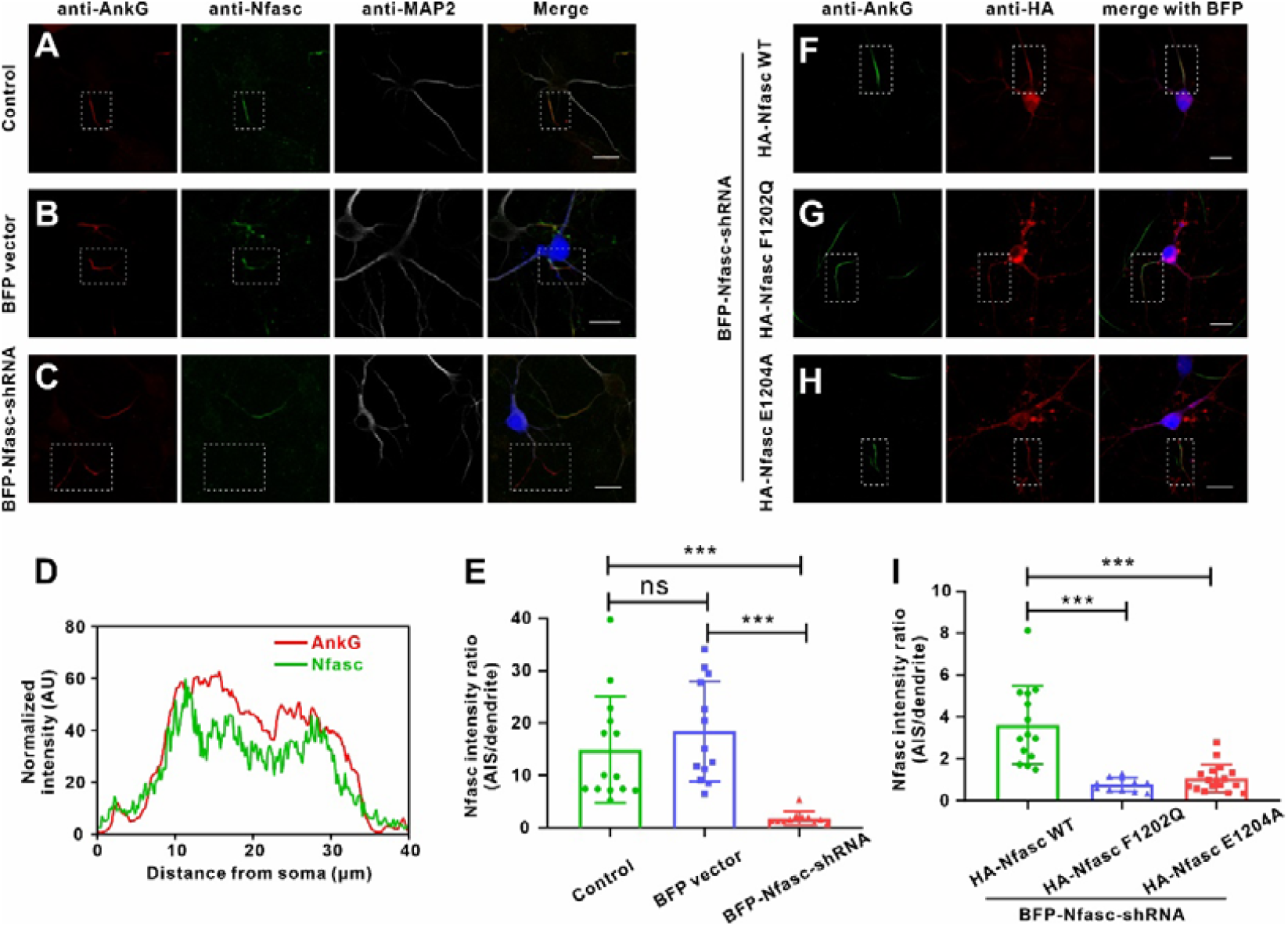
Disruption of AnkG binding abolishes Nfasc enrichment at the AIS. (A-C) WT neurons (A) or neurons transfected with BFP vector (B), or BFP-Nfasc-shRNA (C) and stained for AnkG (red), Nfasc (green), and MAP2 (white). Nfasc-shRNA significantly decreased Nfasc (green) signals in cultured hippocampal neurons. AnkG (red) marks the AIS and MAP2 (white) marks dendrites. The AIS region is marked by dotted boxes. Scale bar: 20 μm. (D) Fluorescence intensity plots of panel A provide a comparison of the immunosignal strength of AnkG (AIS, red) and Nfasc (green), showing the co-localization at the AIS. (E) Quantification of Nfasc fluorescence intensity in Nfasc-shRNA transfected neurons (n=11) compared with the BFP vector transfected neurons (n=13) and WT neurons (n=14). ***p<0.001; ns, not significant. The student’s t-test is performed. Error bars, SEM. (F) The shRNA-resistant HA-Nfasc-WT effectively restored the enrichment at the AIS. BFP marks the Nfasc-shRNA transfected neurons. The AIS region is marked by dotted boxes. Scale bar: 20 μm. (G, H) The HA-Nfasc-F1202Q (G) or HA-Nfasc-E1204A (H) failed to enrich at the AIS. BFP marks the Nfasc-shRNA transfected neurons. The AIS region is marked by dotted boxes. Scale bar: 20 μm. (I) Quantification of Nfasc fluorescence intensity in neurons recused with HA-Nfasc-WT (n=14), HA-Nfasc-F1202Q (n=11), or HA-Nfasc-E1204A (n=17). ***p<0.001; ns, not significant. The student’s t-test is performed. Error bars, SEM.

### L1 Syndrome-associated mutations of L1CAM impair AnkG binding

Nfasc belongs to the L1 superfamily of cell adhesion molecules that includes three other family members, L1CAM, NrCAM, and CHL1 (35). Earlier studies have suggested that all four L1 family members could form complexes with ankyrin family scaffold proteins, most likely through interactions with their respective cytoplasmic tails (15). We confirmed that the proposed Nfasc ABD region in the cytoplasmic tail was highly conserved among L1 members through sequence alignments (Fig. 6A), and further demonstrated through ITC assays that all L1 proteins could bind to AnkG with high affinity (Fig. 6B-D). Interestingly, through investigation of the literatures we found that two mutations (S1226L and Y1231H conversions) in the L1CAM ABD (illustrated in Fig. 6E) have been linked to L1 Syndrome, an inherited mild to severe congenital disorder characterized by corpus callosum hypoplasia, retardation, adducted thumbs, spastic paraplegia and hydrocephalus (36-38).

**Fig. 6.**
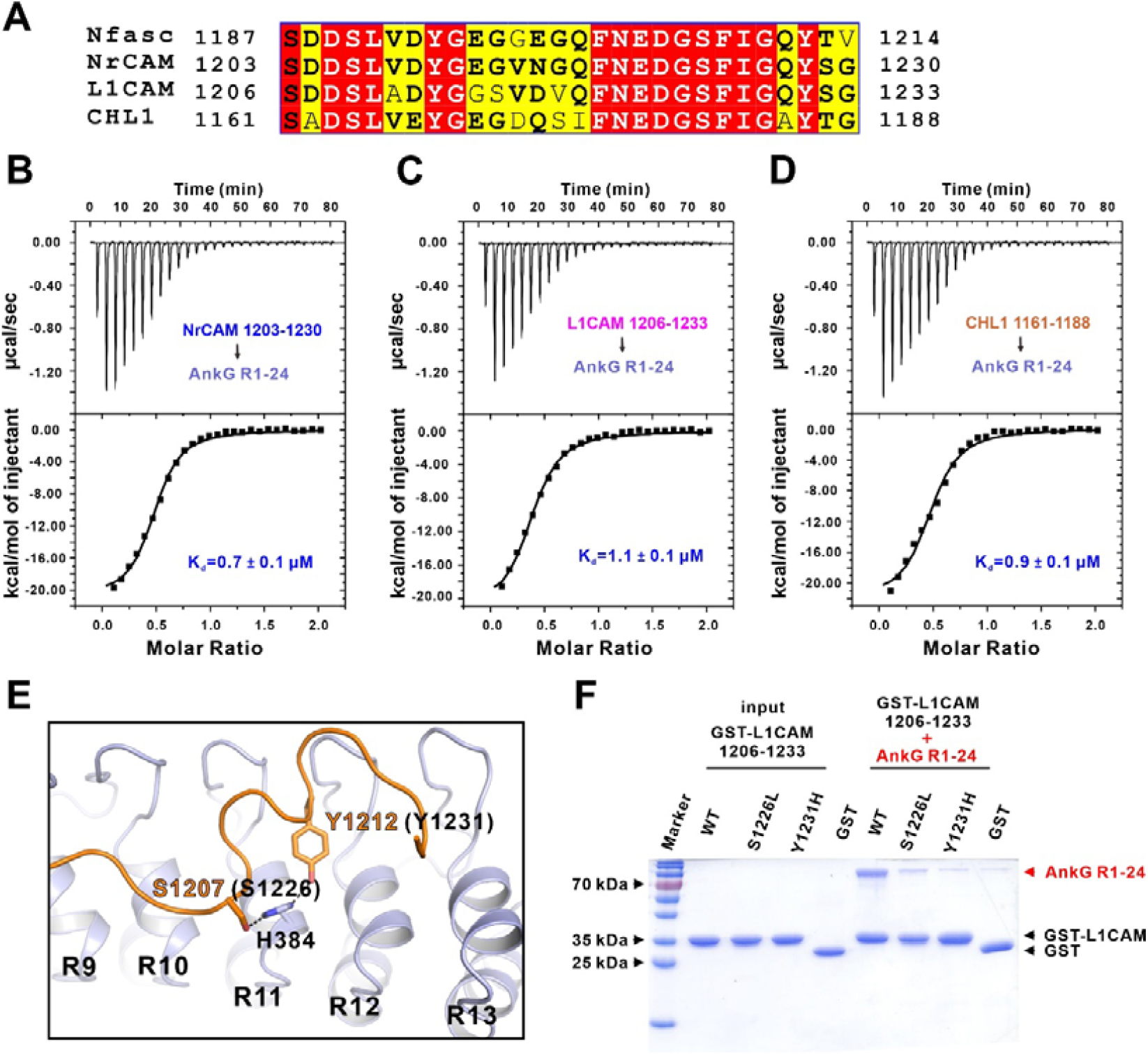
L1 Syndrome-associated mutations of L1CAM impair AnkG binding. (A) Sequence alignment of the ABD region in L1 family members including Nfasc, NrCAM, L1CAM and CHL1. Residues that are identical and highly similar are indicated in read and yeallw boxes, respectively. (B-D) ITC-based measurements of the binding affinity of AnkG R1-24 with NrCAM (B), L1CAM (C), or CHL1 (D), showing all the L1 family proteins can bind to AnkG. (E) Structure model of Nfasc and AnkG showing the residues of Nfasc (S1207 and Y1212) corresponding to the L1 Syndrome-associated residues of L1CAM (S1226 and Y1231). The residue H384 from AnkG is also highlighted in stick model. The hydrogen bonds are shown as dotted lines. (F) GST pull-down assays showing that the S1226L and Y1231H variants of L1CAM significantly decrease the binding with AnkG R1-24.

In our solved crystal structure, the corresponding Ser and Tyr residues of Nfasc form two hydrogen bonds with H384 from AnkG (Fig. 6E). To evaluate whether the two L1 Syndrome-related mutations of L1CAM affect its binding with AnkG, we performed both pull-down and ITC assays to examine their interactions. The results indicated that these two disease-associated mutations indeed resulted in decreased or disrupted binding to AnkG, which could explain the potential pathological mechanism of ankyrin-related trafficking or stabilization of these membrane proteins through formation of membrane microdomain structures (Fig. 6F, & Fig. S6). Taken together, these data suggested that the mode of AnkG-Nfasc interaction revealed in the above data could provide a structural basis for understanding ankyrin-L1 family binding.

## Discussion

The specific localization and molecular composition of the axon initial segment enable its function of initiating action potential and maintaining axonal polarity. AnkG has been proposed to serve as a central coordinator of AIS organization through its capacity to link diverse membrane proteins with cytoskeletal proteins (3, 4, 10, 39). In this study, we systematically characterized the detailed interactions between AnkG and Nfasc by solving the AnkG-Nfasc complex structure and identifying the residues that are essential for binding between Nfasc and AnkG. Moreover, we confirmed Nfasc is enriched at the AIS in a manner dependent on AnkG binding, thus demonstrating a role for Nfasc-AnkG complex formation in maintaining AIS integrity. Finally, we found that mutations in L1CAM linked to L1 Syndrome decrease or disrupt its interactions with AnkG, suggesting that interference with ankyrin-related complex function can contribute to the pathogenesis of neuronal diseases.

Our earlier studies have shown that the N-terminal 24 ANK repeats of ankyrins form an elongated, left-handed helical solenoid structure with the αAs and hairpin loops forming a concave inner groove to coordinate diverse membrane targets binding (24). Furthermore, we demonstrate that AnkG binds to Nfasc through this inner groove of ANK repeats in the present study. Although more than a dozen of membrane binders have been reported to bind with ankyrin family members, the structural information are still very limited. Here, in the solved complex structure, AnkG reserves the typical ANK repeat architecture while Nfasc peptide extends in the elongated inner groove of the ANK repeats, showing the critical role of the inner groove. Interestingly, Nfasc is the first targets proved to interact with ANK repeats in a parallel binding orientation among the reported inner groove-dependent bindings. We further analyzed the amino acids in the inner groove of ANK repeats. Of note, these residues form patterns of preferential distribution to form distinct four layers when the inner groove is critical for targets recognition (Fig. 4). Coordination of these hydrophobic or polar residues layers endows ANK repeat with abilities to bind diverse targets, thus making the ANK repeats an ideal protein-protein interaction module.

Nfasc is essential for the integration of the AIS and thought to be located to the AIS with AnkG at an early stage in the axon differentiation and development period (4, 33, 39, 40). Previous studies have reported that Nfasc is highly mobile when transported to the axonal membrane (30). During moving to the proximal axon via retrograde transport driven by TRIM46-labeled microtubules, Nfasc is retained at the AIS by interact with AnkG (30, 41). The AnkG-Nfasc structure solved in this study provides the structural basis of this crucial complex at the AIS and reveals a new region of Nfasc, which is critical for AnkG interaction besides the FIGQY motif (Fig. 3A). Our biochemical data identified the essential residues (F1202 and E1204) that significantly impact on AnkG binding (Fig. 3B). Moreover, data from cultured hippocampal neurons also confirmed that interference with the AnkG binding in this region of Nfasc failed to rescue Nfasc enrichment at the AIS in neurons (Fig. 5F-I), showing that the AIS localization of Nfasc depends on its binding with AnkG. Interestingly, Nfasc depletion had no obvious influence on AnkG localization at the AIS in the earlier stage (DIV4 to DIV7). However, there are studies reported that shRNA-mediated knockdown of AnkG membrane partners Nav or Nfasc led to a decrease of AnkG concentration and perturbed the AIS formation and maintenance at DIV 14 (33, 42), suggesting that Nfasc indeed plays a vital role in the maintenance of the AIS architecture in a relatively longer period of time in neuronal polarity maintenance.

Previous studies have established that the L1 family proteins function together with ankyrins in diverse membrane microdomains (19). We wonder whether our findings on Nfasc-AnkG interaction could be applicable to other L1 members. Our sequence alignment and ITC data clearly showed that all L1 family proteins could bind to AnkG (Fig. 6A-D). More importantly, we found that two L1 Syndrome associated mutations in L1CAM ABD compromise the binding with AnkG through our structural model and GST pull-down experiments (Fig. 6E, F). In addition, many studies have showed that L1CAM provides linkage to the actin cytoskeleton through AnkB in axons (43-45). Given the high similarity between AnkG and AnkB, we believe that the mutations will also impair L1CAM’s binding to AnkB and our structure may shed light on other functional complex formation beyond the specific AIS region.

In summary, our study establishes the molecular and structural basis for understanding complex formation of AnkG-Nfasc at the AIS, with implications for the maintenance of the AIS integrity and insights into the mechanisms for neuronal diseases.

## Acknowledgments

We thank beamlines BL02U1, BL18U1 and BL19U1 at Shanghai Synchrotron Radiation Facility (SSRF, China) for X-ray beam time. This work was supported by grants from the National Natural Science Foundation of China (22122703, 91953110, 32170767, 31670734, 31971140, 21907033), the Ministry of Science and Technology of the People’s Republic of China (2019YFA0508402), the Fundamental Research Funds for the Central Universities (WK9100000029, WK9100000013), University of Science and Technology of China Research Funds of Double First-Class Initiative (YD9100002006). C.W. is supported by the Chinese Academy of Sciences Pioneer Hundred Talents Program.

## Author contributions

L.H., J.L., and C.W. designed the experiments. L.H. and W.J. performed the experiments. J.L. determined the crystal structure. All authors analyzed the data. L.H., J.L., and C.W. wrote the manuscript. All authors approved the final version of the manuscript. C.W. coordinated the project.

## Data and materials availability

All data needed to evaluate the conclusions in the paper are present in the paper and/or the Supplementary Materials. The atomic coordinates of the AnkG-Nfasc complex have been deposited to the Protein Data Bank under the accession code: 7XCE.

## Materials and Methods

### Constructs, protein expression and purification

All of the protein constructs were cloned into a modified pET32a vector for protein expression and confirmed by DNA sequencing. The fusion constructs used for crystal screening were made by two-step PCR with a linker sequence of “GSLVPRGS” between Nfasc ABD and AnkG ANK repeats. In particular, the consult for solving the crystal structure is made through fusing the Nfasc 1187-1214 (Nfasc ABD) at the N-terminal of AnkG 275-571 (AnkG R8-16) with the above linker sequence. The variant constructs were made by site-directed mutagenesis or standard PCR-based methods and confirmed by DNA sequencing. All the proteins were expressed in BL21 (DE3) *Escherichia coli* cells. The N-terminal thioredxin-His_6_-tagged proteins were purified using Ni-NTA agarose affinity column followed by size-exclusion chromatography (Superdex 200 column, GE Healthcare) in the buffer containing 50 mM Tris-HCl, 1 mM EDTA, 1 mM DTT and 100 mM NaCl or 250 mM NaCl as required at pH 7.8. For crystal screening proteins, the thioredxin-His_6_ tag was removed by incubation with HRV 3C protease at 4°C overnight and separated by size-exclusion chromatography.

### Isothermal titration calorimetry assay

Isothermal titration calorimetry (ITC) measurements were carried out on a VP-ITC MicroCal calorimeter (Malvern) at 25 °C. All proteins were dissolved in the buffer containing 50 mM Tris, 100 mM NaCl, 1 mM EDTA, and 1 mM DTT at pH 7.8. Nfasc proteins (200 μM) were loaded into the syringe, and AnkG proteins (20 μM) were loaded in the cell. Each titration point was obtained by injecting a 10 μl aliquot of syringe protein into the cell at a time interval of 180 sec to ensure that the titration peak returned to the baseline. The titration data were analyzed using the program Origin 7.0 (Microcal) and fitted by the one-site binding model.

### FPLC assay

Analytical gel filtration chromatography was carried out on an AKTA Pure system (GE Healthcare). Proteins were loaded onto a Superose 12 column (GE Healthcare) or a Superdex 200 increase column equilibrated with a buffer containing 50 mM Tris, 100 mM NaCl, 1 mM EDTA, and 1 mM DTT at pH 7.8. All the graphs here were drawn by GraphPad Prism 8.

### GST pull-down assay

For GST pull-down assays, GST-tagged L1CAM 1206-1233 WT, S1226L, and Y1231H variant proteins were purified in the buffer containing 50 mM Tris, 100 mM NaCl, 1 mM EDTA, and 1 mM DTT at pH 7.8, and detected by SDS-PAGE and Coomassie blue staining. The purified AnkG R1-24 proteins (60 μM) were incubated with 20 μM of various GST, GST-L1CAM 1206-1233 WT, GST-L1CAM 1206-1233 S1226L variant, or GST-L1CAM 1206-1233 Y1231H variant for 1 h at 4 °C. The 30 μL GSH-sepharose 4B slurry beads in the protein purified buffer were than incubated with the protein mixture for 30 min at 4 °C. After three times wash with the protein purify buffer, the captured proteins were eluted by 20 μl SDS-PAGE loading dye and detected by coomassie blue staining.

### Protein crystallization and structure determination

Crystals of the AnkG-Nfasc complex were obtained by the hanging drop vapor diffusion method at 16°C under the condition of 25% (w/v) PEP 5/4 PO/OH, 100 mM HEPES (pH 7.5). 25% glycerol was added as the cryo-protectant before diffraction data collection. The diffraction data were collected at Shanghai Synchrotron Radiation Facility at 100 K and processed using HKL3000.

The structure was solved by PHASER software (46) using molecular replacement method with the structure of AnkB R8-14 (PDB: 5Y4E) as the search model. The model of Nfasc was manually built according to the difference electron-density map in COOT (47). Further model modifications and refinements were repeated alternatively using COOT software and PHENIX software. The final model was validated using MolProbity (48) and the statistics are shown in Table S1. The structure figures were made using PyMol software (https://pymol.org/2/).

### Hippocampal neuronal culture and transfection

Primary hippocampal neurons were obtained from newborn C57BL/6 mice hippocampus. The fresh hippocampal tissues were digested with 0.25% trypsin (Life Technologies), and the digestion was terminated by adding 10% FBS (HyClone). The mixture was titrated using a pipette, filtered through a 70 μm sterilized filter, and then centrifuged. The pellet was resuspend gently using DMEM (Life Technologies) added with 1% FBS. Cells were then plated on poly-L-lysine (Sigma-Aldrich) -coated glass coverslips in culture dish at a density of 5 × 10^4^ cells/mL. Neurons were incubated at 37°C in Neurobasal medium (Life Technologies) supplemented with B27, 0.5mM glutamine, 12.5 μM glutamate and 1× Pen Strep and with 5% circulating CO_2_. The medium was changed every 48 h. The shRNA of Nfasc was cloned into a BFP-pll3.7 vector with the sequence 5’-CATCATTCCAACCGTCGTACT-3’ targeting mice Nfasc. The rescue plasmids of Nfasc were cloned into a HA-tagged vector. Hippocampal neurons were transfected by shRNA or rescue plasmids on day 4 using the calcium phosphate-DNA co-precipitation method. On day 7, the neurons were fixed and processed for immunostaining.

### Antibodies and immunofluorescence imaging

Mouse monoclonal antibodies against AnkG (1:1000, N106/36 NeuroMab), rabbit polyclonal antibodies against Neurofascin (1:500, Abcam ab31457), rabbit polyclonal antibodies against HA tag (1:1000, proteintech) and chicken polyclonal antibodies against MAP2 (1:10,000, Abcam) were used in the immunofluorescence imaging. Secondary goat antibodies conjugated to Alexa Fluor 488, 568, or 647 (ThermoFisher) were used at 1:1000 dilutions.

The cultured hippocampal neurons were fixed for 10 min at room temperature with 4% paraformaldehyde (BBI, Shanghai), permeabilized using 0.2 Triton X-100 (BBI, Shanghai) in PBS for 10min, and blocked with blocking buffer (1% BSA, 0.1% Tween 20, BBI, Shanghai). Primary antibodies were diluted in blocking buffer and incubated overnight at 4 °C. The second antibodies were also diluted in blocking buffer and incubated for 1-2 h at room temperature. All cells were washed with PBST (0.1% Tween 20 in PBS) for 3 minutes every time.

All the images in this study were captured using a Zeiss LSM 710 laser-scanning confocal microscope (Zeiss, Germany). The hippocampal neurons were captured using a 63 × 1.4 oil objective. The AIS is defined by the AnkG staining using anti-AnkG antibody. Soma and dendrites were indicated by MAP2 staining using anti-MAP2 antibody. Fluorescence intensity analysis were processed using ImageJ software. The intensity ratios in neurons were quantified and analyzed using GraphPad Prism 8.

## Figure legends

**Fig. S1.**
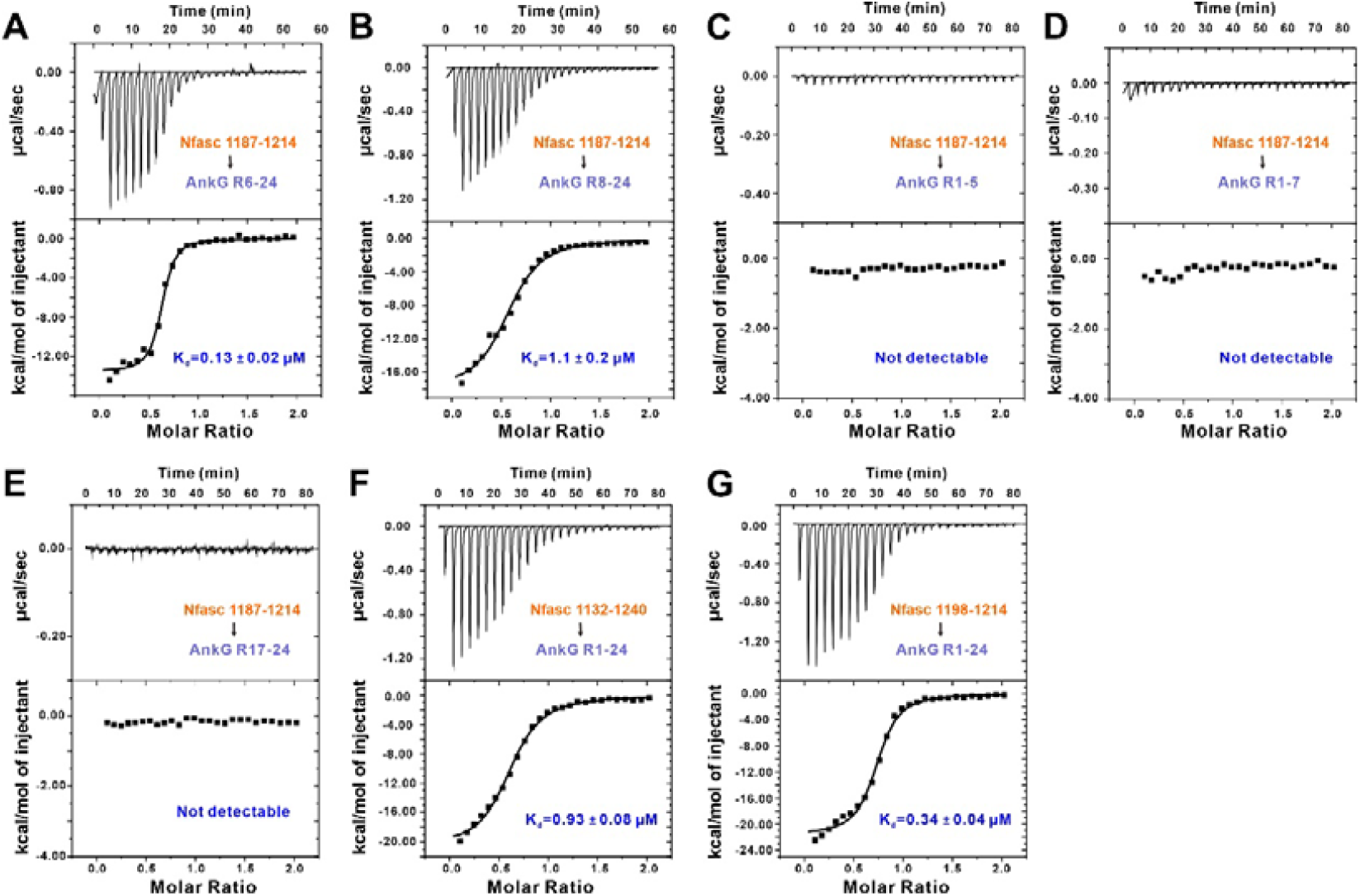
The binding region mapping between AnkG and Nfasc. (A-E) ITC-based measurements of the binding affinities of Nfasc 1187-1214 and various AnkG ANK repeats including R6-24 (A), R8-24 (B), R1-5 (C), R1-7 (D), and R17-24 (E). (F, G) ITC-based measurements of the binding affinities between different Nfasc fragments and AnkG R1-24.

**Fig. S2.**
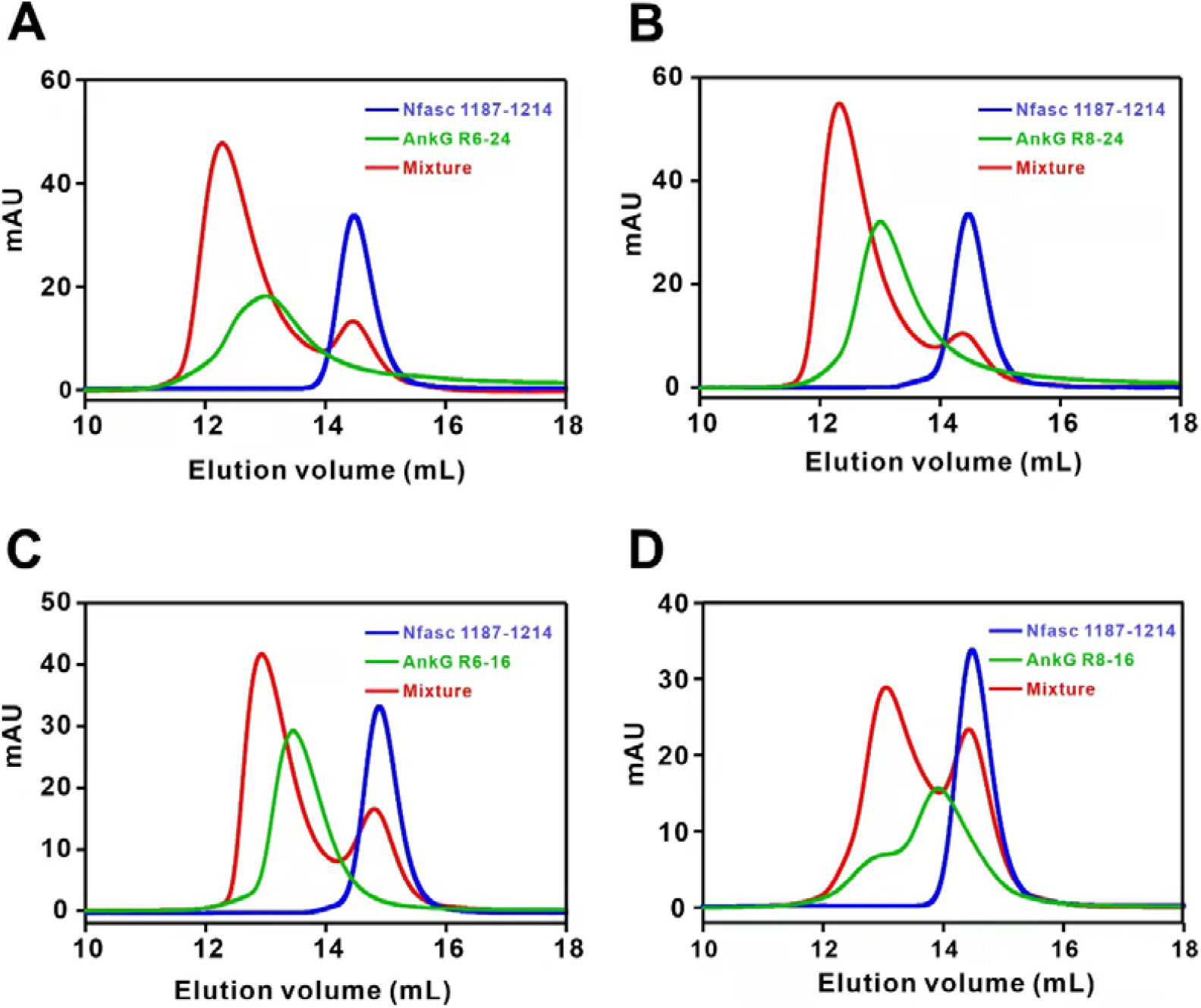
AnkG ANK repeats binds to Nfasc ABD. (A-D) Analytical gel filtration analysis showing that Nfasc ABD (residues 1187-1214) interacts with AnkG ANK repeats R6-24 (A), R8-24 (B), R6-16 (C), and R8-16 (D).

**Fig. S3.**
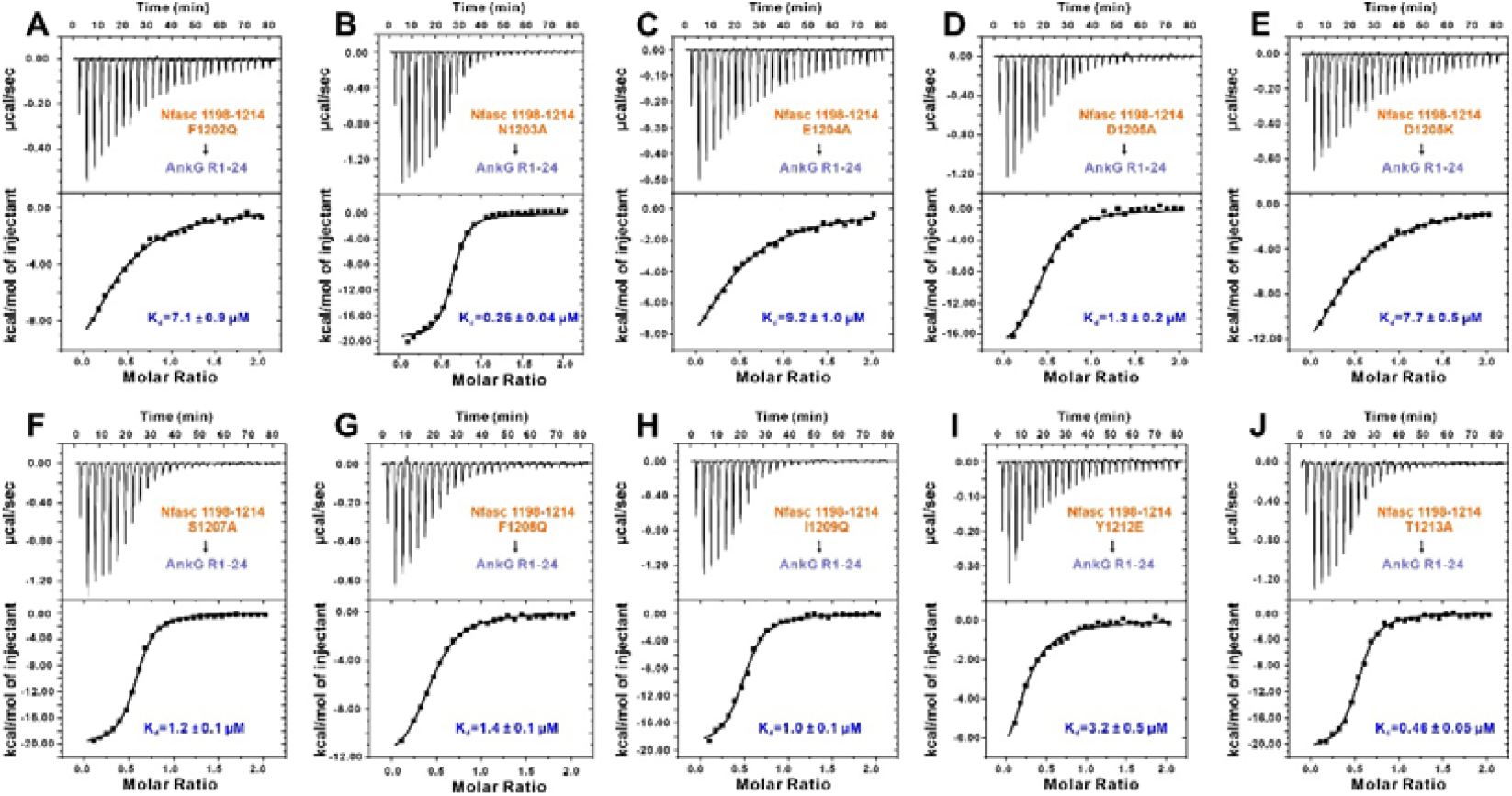
Nfasc mutations decrease the binding affinities between Nfasc and AnkG. ITC-based measurements of the binding affinities of various Nfasc 1198-1214 variants and AnkG R1-24.

**Fig. S4.**
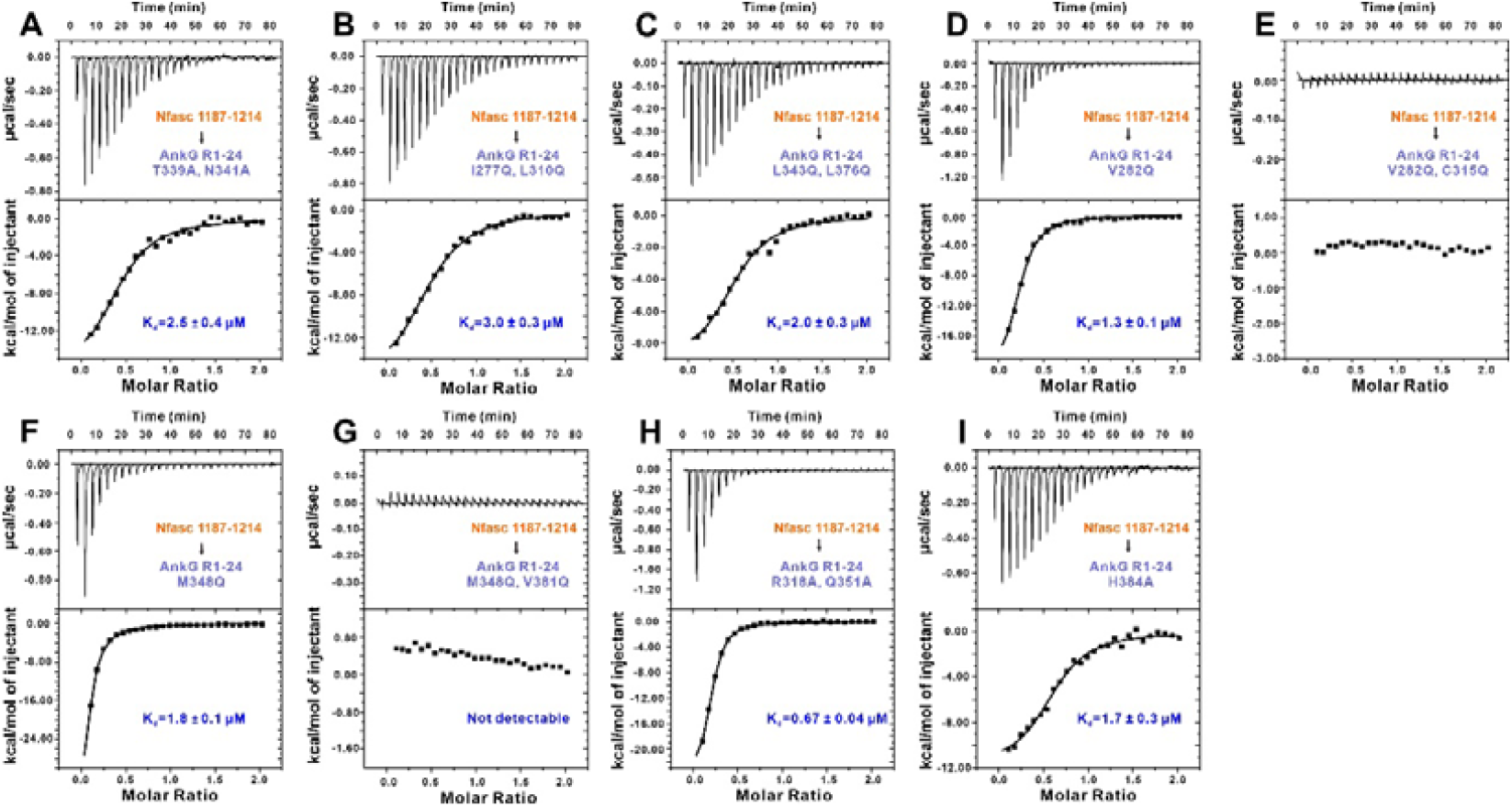
AnkG ANK repeats mutations decrease the binding affinities between Nfasc and AnkG. ITC-based measurements of the binding affinities of Nfasc 1187-1214 and various AnkG R1-24 variants.

**Fig. S5.**
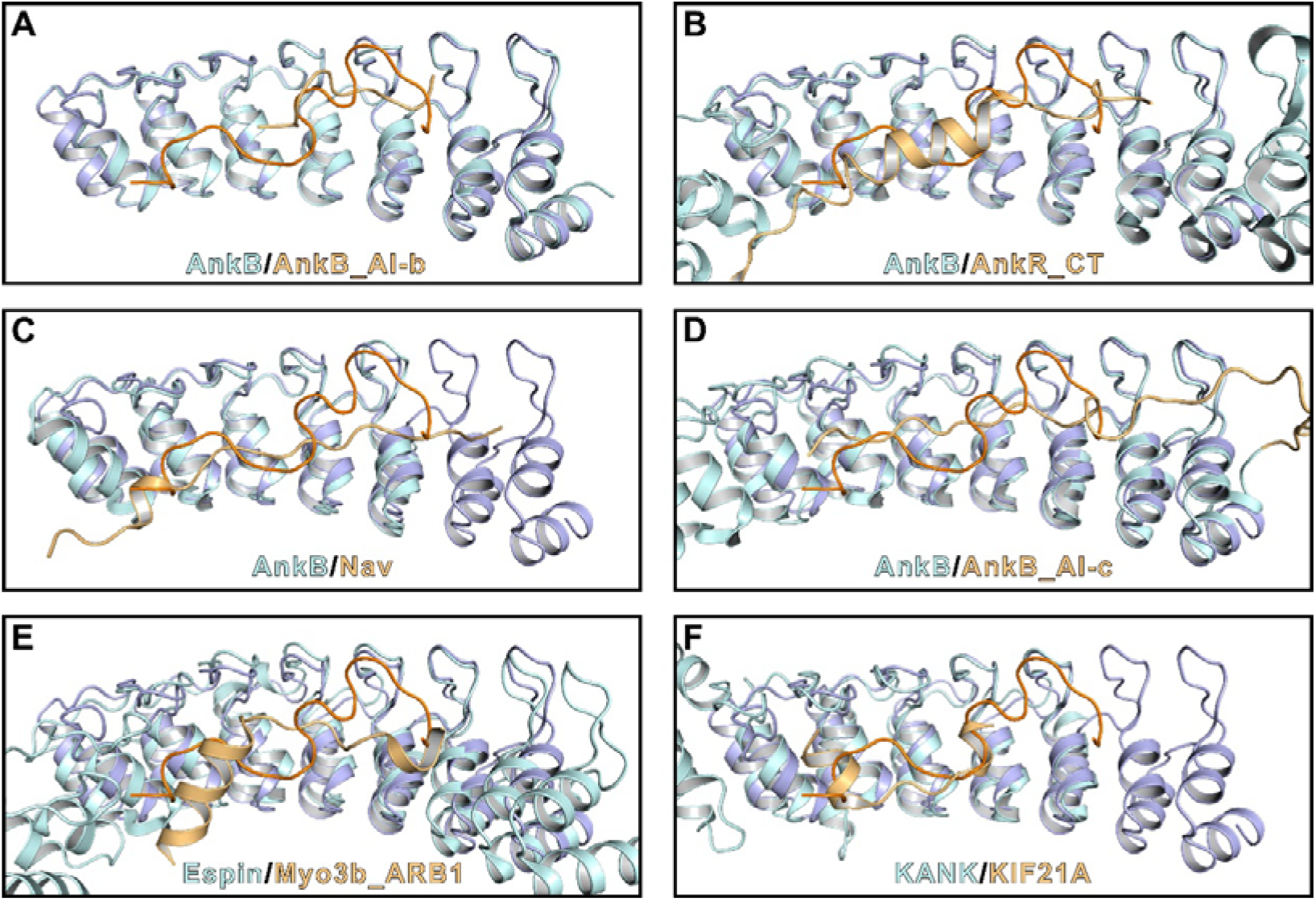
Structure alignments of AnkG-Nfasc complex with other ANK repeats containing proteins in complex with respective cytoplasmic peptides. (A) Structure alignment of AnkG-Nfasc complex and AnkB/AnkB AI-b complex, PDB ID: 5Y4E. (B) Structure alignment of AnkG-Nfasc complex and AnkB/AnkR_CT complex, PDB ID: 4RLV. (C) Structure alignment of AnkG-Nfasc complex and AnkB/Nav1.2 complex, PDB ID: 4RLY. (D) Structure alignment of AnkG-Nfasc complex and AnkB/AnkB AI-c complex, PDB ID: 5Y4F. (E) Structure alignment of AnkG-Nfasc complex and Espin/Myo3b_ARB1 complex, PDB ID: 5ET1. (F) Structure alignment of AnkG-Nfasc complex and KANK1/KIF21A complex, PDB ID: 5YAY. The color schemes are in line with the corresponding proteins in each individual panel.

**Fig. S6.**
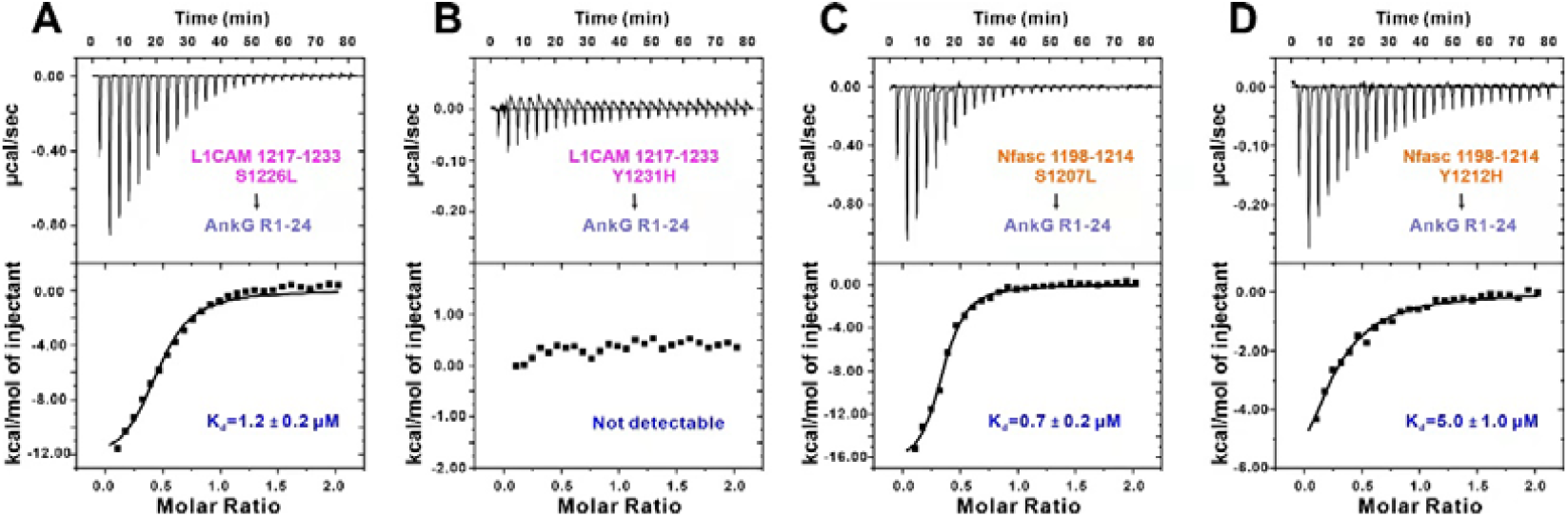
L1 Syndrome-associated mutations decrease the binding affinities between L1CAM and AnkG. (A, B) ITC-based measurements of the binding affinities of between L1CAM 1217-1233 S1226L (A) or L1CAM 1217-1233 Y1231H (B) and AnkG R1-24. (C, D) ITC-based measurements of the binding affinities of between Nfasc 1198-1214 S1207L (C) or Nfasc 1198-1214 Y1212H (D) and AnkG R1-24.

## Tables

**Table S1.** Statistics of X-ray Crystallographic Data Collection and Model refinement.

| **Data collection** |  |
| --- | --- |
| Data sets | AnkG/Nfasc |
| Space group | *P4<sub>1</sub>2<sub>1</sub>1* |
| Wavelength (Å) | 0.97915 |
| Unit Cell Parameters (Å) | a=b=97.877, c=89.101 |
| | $\alpha=\beta=\gamma=90^\circ$ |
| Resolution range (Å) | 50-2.50 (2.54-2.50) |
| No. of unique reflections | 15440 (760) |
| Redundancy | 5.4 (5.6) |
| I/$\sigma$ | 21.7 (2.7) |
| Completeness (%) | 99.6 (99.9) |
| R<sub>merge</sub><sup>a</sup> (%) | 12.4 (91.3) |
| CC<sub>1/2</sub> (last resolution shell)<sup>b</sup> | 0.786 |
| **Structure refinement** |  |
| Resolution (Å) | 50-2.50 (2.59-2.50) |
| R<sub>cryst</sub><sup>c</sup>/R<sub>free</sub><sup>d</sup> (%) | 19.10/23.70 (25.94/31.69) |
| rmsd bonds (Å) / angles (°) | 0.008 / 1.018 |
| Average B factor (Å<sup>2</sup>)<sup>e</sup> | 51.6 |
| No. of atoms |  |
| Protein atoms | 1800 |
| Water | 30 |
| Ligands | 11 |
| No. of reflections |  |
| Working set | 14671 (1427) |
| Test set | 744 (72) |
| Ramachandran plot regions<sup>d</sup> |  |
| Favored (%) | 98.77 |
| Allowed (%) | 1.23 |
| Outliers (%) | 0 |

Numbers in parentheses represent the value for the highest resolution shell.
a. $R_{\text{merge}} = \sum |I_i - \langle I \rangle| / \sum I_i$ , where $I_i$ is the intensity of measured reflection and $\langle I \rangle$ is the mean intensity of all symmetry-related reflections.
- b. $CC_{1/2}$ were defined by Karplus and Diederichs (49). - c. $R_{\text{cryst}} = \sum ||F_{\text{calc}}| - |F_{\text{obs}}|| / \sum F_{\text{obs}}$ , where $F_{\text{obs}}$ and $F_{\text{calc}}$ are observed and calculated structure factors. - d. $R_{\text{free}} = \sum_T ||F_{\text{calc}}| - |F_{\text{obs}}|| / \sum F_{\text{obs}}$ , where T is a test data set of about 5% of the total unique reflections randomly chosen and set aside prior to refinement. - e. B factors and Ramachandran plot statistics are calculated using MOLPROBITY (50).

## References

1. Kole MH & Stuart GJ (2012) Signal processing in the axon initial segment. Neuron 73(2):235–247.

2. Zhou D, et al. (1998) AnkyrinG is required for clustering of voltage-gated Na channels at axon initial segments and for normal action potential firing. J Cell Biol 143(5):1295–1304.

3. Hedstrom KL, Ogawa Y, & Rasband MN (2008) AnkyrinG is required for maintenance of the axon initial segment and neuronal polarity. J Cell Biol 183(4):635–640.

4. Rasband MN (2010) The axon initial segment and the maintenance of neuronal polarity. Nat Rev Neurosci 11(8):552–562.

5. Leterrier C (2018) The Axon Initial Segment: An Updated Viewpoint. J Neurosci 38(9):2135–2145.

6. Albrecht D, et al. (2016) Nanoscopic compartmentalization of membrane protein motion at the axon initial segment. J Cell Biol 215(1):37–46.

7. Quistgaard EM, Nissen JD, Hansen S, & Nissen P (2021) Mind the Gap: Molecular Architecture of the Axon Initial Segment - From Fold Prediction to a Mechanistic Model of Function? J Mol Biol 433(20):167176.

8. Jenkins SM & Bennett V (2001) Ankyrin-G coordinates assembly of the spectrin-based membrane skeleton, voltage-gated sodium channels, and L1 CAMs at Purkinje neuron initial segments. J Cell Biol 155(5):739–746.

9. Jones SL & Svitkina TM (2016) Axon Initial Segment Cytoskeleton: Architecture, Development, and Role in Neuron Polarity. Neural Plast 2016:6808293.

10. Huang CY & Rasband MN (2018) Axon initial segments: structure, function, and disease. Ann N Y Acad Sci 1420(1):46–61.

11. Kole MH, et al. (2008) Action potential generation requires a high sodium channel density in the axon initial segment. Nat Neurosci 11(2):178–186.

12. Kole MH & Stuart GJ (2008) Is action potential threshold lowest in the axon? Nat Neurosci 11(11):1253–1255.

13. Kuijpers M, et al. (2016) Dynein Regulator NDEL1 Controls Polarized Cargo Transport at the Axon Initial Segment. Neuron 89(3):461–471.

14. Ye J, et al. (2020) Mechanistic insights into the interactions of dynein regulator Ndel1 with neuronal ankyrins and implications in polarity maintenance. Proc Natl Acad Sci U S A 117(2):1207–1215.

15. Davis JQ & Bennett V (1994) Ankyrin binding activity shared by the neurofascin/L1/NrCAM family of nervous system cell adhesion molecules. J Biol Chem 269(44):27163–27166.

16. Koticha D, et al. (2006) Neurofascin interactions play a critical role in clustering sodium channels, ankyrin G and beta IV spectrin at peripheral nodes of Ranvier. Dev Biol 293(1):1–12.

17. Sobotzik JM, et al. (2009) AnkyrinG is required to maintain axo-dendritic polarity in vivo. Proc Natl Acad Sci U S A 106(41):17564–17569.

18. Davis LH, Davis JQ, & Bennett V (1992) Ankyrin regulation: an alternatively spliced segment of the regulatory domain functions as an intramolecular modulator. J Biol Chem 267(26):18966–18972.

19. Bennett V & Lorenzo DN (2013) Spectrin- and ankyrin-based membrane domains and the evolution of vertebrates. Curr Top Membr 72:1–37.

20. Kordeli E, Lambert S, & Bennett V (1995) AnkyrinG. A new ankyrin gene with neural-specific isoforms localized at the axonal initial segment and node of Ranvier. J Biol Chem 270(5):2352–2359.

21. Jenkins PM, et al. (2015) Giant ankyrin-G: a critical innovation in vertebrate evolution of fast and integrated neuronal signaling. Proc Natl Acad Sci U S A 112(4):957–964.

22. Garrido JJ, et al. (2003) A targeting motif involved in sodium channel clustering at the axonal initial segment. Science 300(5628):2091–2094.

23. Pan Z, et al. (2006) A common ankyrin-G-based mechanism retains KCNQ and NaV channels at electrically active domains of the axon. J Neurosci 26(10):2599–2613.

24. Wang C, et al. (2014) Structural basis of diverse membrane target recognitions by ankyrins. Elife 3.

25. Chen K, Li J, Wang C, Wei Z, & Zhang M (2017) Autoinhibition of ankyrin-B/G membrane target bindings by intrinsically disordered segments from the tail regions. Elife 6.

26. Hortsch M (1996) The L1 family of neural cell adhesion molecules: old proteins performing new tricks. Neuron 17(4):587–593.

27. Herron LR, Hill M, Davey F, & Gunn-Moore FJ (2009) The intracellular interactions of the L1 family of cell adhesion molecules. Biochem J 419(3):519–531.

28. Kriebel M, Wuchter J, Trinks S, & Volkmer H (2012) Neurofascin: a switch between neuronal plasticity and stability. Int J Biochem Cell Biol 44(5):694–697.

29. Zonta B, et al. (2011) A critical role for Neurofascin in regulating action potential initiation through maintenance of the axon initial segment. Neuron 69(5):945–956.

30. Ghosh A, Malavasi EL, Sherman DL, & Brophy PJ (2020) Neurofascin and Kv7.3 are delivered to somatic and axon terminal surface membranes en route to the axon initial segment. Elife 9.

31. Tuvia S, Garver TD, & Bennett V (1997) The phosphorylation state of the FIGQY tyrosine of neurofascin determines ankyrin-binding activity and patterns of cell segregation. Proc Natl Acad Sci U S A 94(24):12957–12962.

32. Jenkins SM, et al. (2001) FIGQY phosphorylation defines discrete populations of L1 cell adhesion molecules at sites of cell-cell contact and in migrating neurons. J Cell Sci 114(Pt 21):3823–3835.

33. Alpizar SA, Baker AL, Gulledge AT, & Hoppa MB (2019) Loss of Neurofascin-186 Disrupts Alignment of AnkyrinG Relative to Its Binding Partners in the Axon Initial Segment. Front Cell Neurosci 13:1.

34. Hedstrom KL, et al. (2007) Neurofascin assembles a specialized extracellular matrix at the axon initial segment. J Cell Biol 178(5):875–886.

35. Hortsch M (2000) Structural and functional evolution of the L1 family: are four adhesion molecules better than one? Mol Cell Neurosci 15(1):1–10.

36. Fransen E, et al. (1995) CRASH syndrome: clinical spectrum of corpus callosum hypoplasia, retardation, adducted thumbs, spastic paraparesis and hydrocephalus due to mutations in one single gene, L1. Eur J Hum Genet 3(5):273–284.

37. Yamasaki M, Thompson P, & Lemmon V (1997) CRASH syndrome: mutations in L1CAM correlate with severity of the disease. Neuropediatrics 28(3):175–178.

38. Sztriha L, Frossard P, Hofstra RM, Verlind E, & Nork M (2000) Novel missense mutation in the L1 gene in a child with corpus callosum agenesis, retardation, adducted thumbs, spastic paraparesis, and hydrocephalus. J Child Neurol 15(4):239–243.

39. Leterrier C (2016) The Axon Initial Segment, 50Years Later: A Nexus for Neuronal Organization and Function. Curr Top Membr 77:185-233.

40. Torii T, et al. (2020) NuMA1 promotes axon initial segment assembly through inhibition of endocytosis. J Cell Biol 219(2).

41. Fréal A, et al. (2019) Feedback-Driven Assembly of the Axon Initial Segment. Neuron 104(2):305-321.e308.

42. Leterrier C, et al. (2017) Ankyrin G Membrane Partners Drive the Establishment and Maintenance of the Axon Initial Segment. Front Cell Neurosci 11:6.

43. Nishimura K, et al. (2003) L1-dependent neuritogenesis involves ankyrinB that mediates L1-CAM coupling with retrograde actin flow. J Cell Biol 163(5):1077–1088.

44. Buhusi M, Schlatter MC, Demyanenko GP, Thresher R, & Maness PF (2008) L1 interaction with ankyrin regulates mediolateral topography in the retinocollicular projection. J Neurosci 28(1):177–188.

45. Yang R, et al. (2019) ANK2 autism mutation targeting giant ankyrin-B promotes axon branching and ectopic connectivity. Proc Natl Acad Sci U S A 116(30):15262–15271.

46. Mccoy AJ, et al. (2007) Phaser crystallographic software. J. Appl. Crystallogr. 40:658-674.

47. Emsley P, Lohkamp B, Scott WG, & Cowtan K (2010) Features and development of Coot. Acta Crystallogr. D Biol. Crystallogr. 66:486–501.

48. Chen VB, et al. (2010) MolProbity: all-atom structure validation for macromolecular crystallography. Acta Crystallogr. D Biol. Crystallogr. 66:12–21.

49. Karplus PA & Diederichs K (2012) Linking crystallographic model and data quality. Science 336(6084):1030–1033.

50. Chen VB, et al. (2010) MolProbity: all-atom structure validation for macromolecular crystallography. Acta Crystallographica. Section D, Biological Crystallography 66(Pt 1):12–21.

